# The claustrum is critical for maintaining working memory information

**DOI:** 10.1101/2024.10.28.620649

**Authors:** Anindya S. Bhattacharjee, Chieko Huber, Burak Unsal, Jean-Rodolphe Renfer, Ivan Rodriguez, Alan Carleton

**Author notes:** Correspondence should be addressed to A.C. or I.R. Senior authors contributed equally to the work (Ivan Rodriguez, Alan Carleton).

## Abstract

Working memory (WM) enables the mammalian brain to temporarily store and manipulate information, supporting cognitive tasks and communication processes^1,2^. Rather than depending on a single specialized area, WM is thought to operate through a distributed network spanning cortical and subcortical regions^3–5^. A dedicated WM storage area would likely require broad reciprocal connections with various cortical regions to accommodate the diverse range of information WM retains. The claustrum (CLA), with its extensive bidirectional connections to the neocortex^6–9^, presents a compelling candidate for such a role. Here, we examined the involvement of the CLA in WM processes by recording CLA neuronal activity in mice engaged in olfactory and tactospatial delayed non-match-to-sample WM tasks. We identified cue-selective and delay-specific neurons in the CLA that maintained activity for tens of seconds after the stimulus presentation ended. Additionally, population activity in the CLA allowed for decoding of cue identity post-stimulus, although this signal gradually declined over time, aligning with animal behavior. Remarkably, both chemo- and optogenetic inhibition of CLA neurons severely impaired WM performance across multiple types of stored information, highlighting the CLA’s critical role during both cue encoding, delay periods, and target comparison phases. These findings challenge the view that no single brain area is essential for WM storage and support a role for the CLA as an essential WM storage hub.

## Main text

Working memory (WM) is a fundamental cognitive function that temporarily stores information required for complex processes such as reasoning, learning, communication, and decision-making. The prefrontal cortex (PFC), particularly the dorsolateral PFC in primates and its equivalent, the medial PFC in rodents^10^, is widely recognized as a core brain area for WM function^3,11,12^. Initially, the neural correlate of WM was thought to be represented in the PFC’s sustained firing of pyramidal neurons during delay periods of response tasks^13–19^. However, as recording technology advanced and allowed for larger groups of neurons to be monitored, transient rather than sustained responses became more commonly observed, with population activity sometimes disappearing during the delay only to reemerge later^12,20–23^. These findings led to theories that information might be maintained not solely through PFC activity but also through short-term plasticity in synaptic connections during silent intervals^24,25^ or through a combination of both mechanisms^26^.

Although contributing to WM, the PFC is not the sole region involved, and evidence suggests it may play a more prominent role during WM task learning than in information storage after the task is mastered^18^. Emerging research in humans and other mammals indicates that WM relies on a distributed network, encompassing sensory cortices^27–30^, associative areas within the frontal^13–19,31,32^ and parietal^33,34^ lobes, the basal ganglia^35,36^, thalamus^37–39^, and hippocampus^40–42^. This organization supports the integration of specialized information across brain regions: the frontal cortex helps organize task-relevant details, sensory and parietal cortices store sensory and spatial information during short delays. The networked nature of WM allows for flexible, adaptive functioning that remains resilient to disruptions within individual network components, with higher cortical areas increasingly contributing as delay periods extend^32^.

The distributed nature of WM invites the question of how storage across multiple regions is coordinated. Conceptually, one or a few brain regions could temporarily consolidate sensory inputs, creating a unified platform to transform this information into appropriate behavioral responses^43,44^. Investigating the presence of such a storage hub could be crucial for understanding how the brain integrates sensory information to guide action. The claustrum (CLA), a subcortical nucleus situated between the striatum and insular cortex, may serve this function, as it predominantly consists of glutamatergic projection neurons and is extensively reciprocally connected to the neocortex^6–9,45^. CLA inputs can modulate cortical circuit activity in complex ways that depend on stimulation duration, cortical regions, and cell types, as shown in passive mice using optogenetic stimulation^45–47^. Given these unique features, we tested whether the CLA might function as a localized storage center, capable of maintaining diverse information types in WM.

### Working memory performance across distinct sensory modalities in mice

We first aimed to compare WM performance across two different sensory dimensions, olfactory and tactospatial information, in head-restrained mice (**Fig. 1a**). To achieve this goal, we employed a go/no-go paradigm, which entailed presenting sequences of either two odorants or two tactile stimuli as sensory cues. A sample cue was presented for one second and was required to be retained during a delay period. Subsequently, the mouse encountered a target cue, which could either match the sample stimulus (match trials) or be a different cue (non-match trials). To earn a reward, mice were instructed to lick a reward tube within a defined response window, specifically for non-match trials, thus following a delayed non-match-to-sample design. Successful trials demonstrated the ability of the animals to transiently store information in their WM for future comparison (**Fig. 1a**), as neither the first nor the second cue alone could reliably predict the reward outcome. Pairs of cues for olfactory WM (OWM) and tactospatial WM (TSWM) tasks were defined using two easily discriminated odorants^48^ and an airflow stimulation of the left or right whisker pad, respectively (**Fig. 1a**). We computed the percentage of correct responses, or accuracy, specifically referring to trials in which a mouse licked for non-match trials and refrained from licking for match trials (responses to the contrary were considered errors). Furthermore, we distinguished between false alarms, correct rejections, hits, and misses to calculate the sensitivity index, D-prime. The mice were initially trained to perform an OWM task with a fixed intercue delay of five seconds. They rapidly learned the task and achieved higher performance levels (**Fig. 1b,c**). Subsequently, they underwent multiple daily sessions during which the intercue delay was randomly varied across trials. As anticipated for a WM task, performance declined as the intercue delay increased (**Fig. 1d**). Next, the mice were trained to perform the TSWM task with a fixed intercue delay of one second. They swiftly adjusted to the new sensory dimension and attained high-performance levels (**Fig. 1b,c**). Finally, the animals underwent sessions during which the tactospatial intercue delay was randomly adjusted across trials. At both population average and individual levels of analysis, WM performance showed an exponential decline with increasing intercue delay for both tasks (**Fig. 1d,e**). While the performance consistently appeared to be slightly greater for the OWM task than for the TSWM task (**Fig. 1d,f,g**), the memory half-life remained independent of the sensory dimension employed (∼20 and 30 seconds for the median and mean, respectively; **Fig. 1h,i**). In summary, the assessment of WM performance across different sensory dimensions can reliably be conducted in mice.

**Figure 1.**
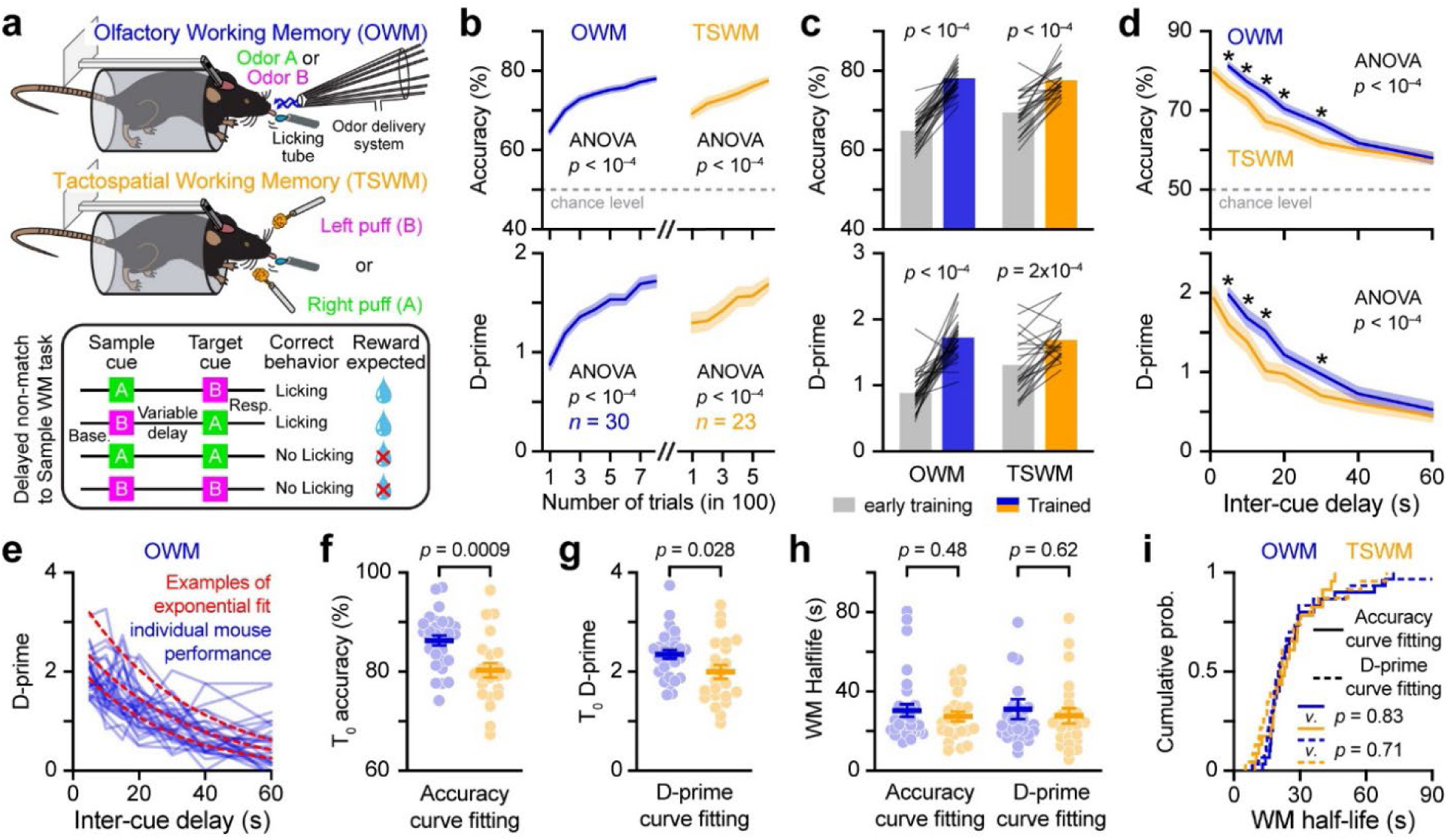
Working memory performance across distinct sensory modalities in mice. **a,** Schematic representation of the behavioral paradigm designed to evaluate olfactory and tactospatial WM (OWM and TSWM, respectively) in head-restrained mice. For each trial, both sample and target cues were presented for one second (see main text and methods for further details). **b,** Learning curves for OWM and TSWM tasks (fixed intercue delay of 5 and 1s, respectively; the number of mice is indicated; one-way repeated measures ANOVA). **c,** Change in performance between the first and the last 100 trials of the training phase shown in **b** for each mouse (black lines; paired t-test) and the mean population performance (colored bars). **d,** WM performances plotted as a function of the intercue delay duration (2WRM ANOVA, post-hoc Fisher’s test *: P < 0.05). **e,** Variability of WM performance across all individual animals shown for the D-prime parameter computed during the OWM task. Examples of exponential fit (used to calculate the amplitude at 0 delay and the half-life) of a few individual performance curves are illustrated (red dashed lines). **f-h,** Comparison of WM performance amplitude at 0 delay and half-life for OWM and TSWM tasks (unpaired t-test). **i,** Cumulative probability distribution of half-lives for OWM and TSWM tasks (Kolmogorov-Smirnov test). Data in **b**, **d**, **f-h** are presented as mean ± SEM. See **Supplementary Table 1** for detailed statistics.

### Claustrum ensembles maintain cue information during working memory

We then proceeded to monitor the activity of genetically identified CLA neurons using microendoscopic calcium imaging in mice executing the WM tasks described above (**Fig. 2a**). We achieved this by virally expressing the genetically encoded calcium indicator GCaMP6f in CLA projection neurons expressing the vesicular glutamate transporter 2 (VGLUT2), a genetic marker of these neurons^45^. Changes in calcium activity were tracked as an indicator of modulation in firing rates. To mitigate the risk of phototoxicity, imaging was restricted to trials with delays of 5, 15, and 30 seconds. Some CLA neurons reliably altered their activity across trials during the WM tasks and could be categorized into three groups (**Fig. 2b-e**). The first type, referred to as stimulus neurons, displayed increased activity immediately upon presentation of the sample cue. These neurons exhibited complex response patterns during the delay period, with some periods of inhibition and/or excitation. The second group, termed delay neurons, showed heightened calcium activity specifically during the intercue delay. Interestingly, numerous delay neurons exhibited activity consistently observed at specific time intervals during the delay across various trials, reminiscent of the time cells previously documented in the hippocampus^40,41,49–51^ and entorhinal cortex^52^. Finally, a third group exhibited inhibition during extensive portions of the trial.

**Figure 2.**
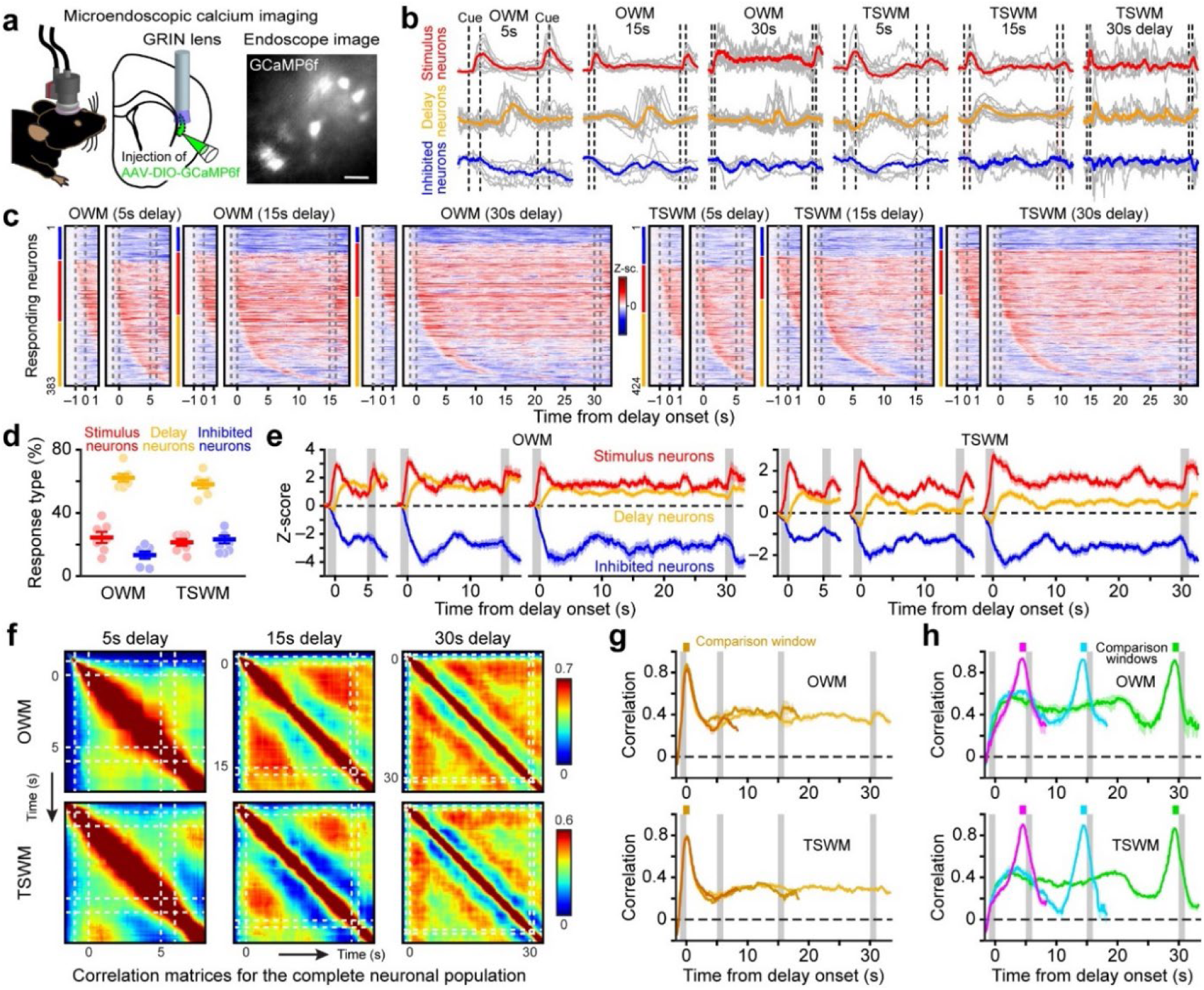
Persistence of claustrum neuron responses after sensory cue presentation. **a,** Schema of the calcium imaging procedure. Image of CLA neurons expressing the genetically encoded calcium indicator GCaMP6f captured through the GRIN lens of the endoscope camera (scale bar: 50 µm). **b,** Examples of individual neuron responses during OWM and TSWM tasks. Grey and colored traces represent individual and averaged correct trials for different response types, respectively. Dashed black lines mark cue application periods. **c,** Pseudo-colored heat maps presenting trial-averaged responses from correct trials for all stimulus, delay, and inhibited neurons (distribution indicated by the color lines on the left, same code as in **b**). Neurons were pooled from 7 mice and sorted by response type and latency (z-score min: -15, max: 20 and 15 for OSW and TSWM maps, respectively). For each delay duration, an expanded zoom including the first second of the delay is shown. Dashed grey lines indicate cue applications. **d,** Quantification of the different response types recorded in different mice (each data point represents the mean value computed across different delays for each animal; data presented as mean ± SEM). **e,** Average population calcium activity (data presented as mean ± SD). Grey bars represent cue applications. **f,** Correlation matrices showing all possible correlations computed between pairs of population activity vectors at different time points (time bin = 100 ms). Dashed white lines represent cue application. **g-h,** Evolution of ensemble correlation over time computed for different trial structures (sample A-target B, sample A-target A, sample B-target A, sample B-target B; data are presented as mean ± SD) and plotted for various delay durations during the two WM tasks. The comparison windows (1s long) are color-coded and indicated on top of the graphs. Grey bars represent cue applications. See **Supplementary Table 1** for detailed statistics.

Notably, we observed these distinct response types regardless of the sensory dimensions investigated, the delay duration, or the individual animal (**Fig. 2b-d**). On average, ∼25% of the cells were categorized as stimulus neurons and ∼60% as delay neurons (**Fig. 2d**). The average population responses exhibited sustained changes in activity until the target cue was presented for all three groups (**Fig. 2e**). While individual neurons seldom displayed sustained activity throughout both the cue and delay periods, the overall CLA population responded to the sample cue and maintained changes in activity throughout the delay, even for durations as long as 30 seconds (**Fig. 2c,e**). To further investigate the dynamics of population responses across different delay durations or sensory dimensions, we constructed a population vector with N dimensions, where N represents the total number of responding neurons recorded in all animals. Within each dimension, we calculated the mean calcium z-score amplitude averaged across all correct trials for all task-delay pairs every 100 ms. Consequently, the temporal evolution of CLA population activity was characterized by vector time series for each trial duration and WM task. One approach to capturing the temporal evolution of population activity during a trial was to compute the similarity between time series vectors using the Pearson correlation coefficient (**Fig. 2f-h**). The systematic correlation analysis across all possible time windows is depicted in correlation matrices (**Fig. 2f**). The presentation of the sample cue triggered a population response that persisted throughout the delay period, although correlation values varied over time, indicating fluctuations in the ensemble of active neurons. Comparing all vectors to those recorded at the peak response time of stimulus neurons revealed a gradual decline in correlation during the delay before reaching a plateau (**Fig. 2g**). Nevertheless, the correlation remained consistently above baseline values, suggesting that a portion of the neuronal population sustained an activity pattern initiated by the sample cue. This was further supported when comparison windows were aligned with the very end of each delay period, as correlations during the cue presentation window again exceeded baseline levels. Intriguingly, we observed more pronounced fluctuations in correlation values over time in the latter analysis, indicating that representation underwent greater ensemble reorganization during the delay (**Fig. 2h**). In summary, regardless of the delay duration and the sensory domain studied, sensory cues elicited changes in CLA neuron activity that persisted throughout the entire delay period, albeit with evolving population dynamics over time.

Does the CLA retain information about the identity of the sample cue and sustain it during the delay? To investigate these questions, our initial focus was identifying cue-selective responses within the CLA population for each WM task. We observed that neurons exhibited a selective increase in calcium activity during the presentation of one cue but not the other (**Fig. 3a**). Intriguingly, the response to the alternative cue could sometimes elicit either an inhibitory response or an increased calcium activity during the delay (**Fig. 3a,b**). This finding suggested that the three previously identified groups of neurons might reflect functional tuning properties for different cues rather than distinct neuronal subpopulations^53,54^. Subsequently, we examined the activity dynamics of the entire CLA population elicited by the two cues of each sensory dimension. Employing a dimensionality reduction algorithm, we plotted the evolution of the population activity during the averaged correct trials in the space defined by the three principal components carrying the most variance. Upon cue presentation, the population activity transitioned from the baseline state and initiated a cue-specific trajectory that remained distinct from the alternative cue-evoked representation not only during the cue presentation but also throughout the delay period (**Fig. 3c**). These observations were consistent across delay durations and sensory dimensions probed. As anticipated from the correlation analyses (**Fig. 2f-h**), the representation changed during the delay period but remained sufficiently different to reflect the identity of the presented cue (**Fig. 3c**).

**Figure 3.**
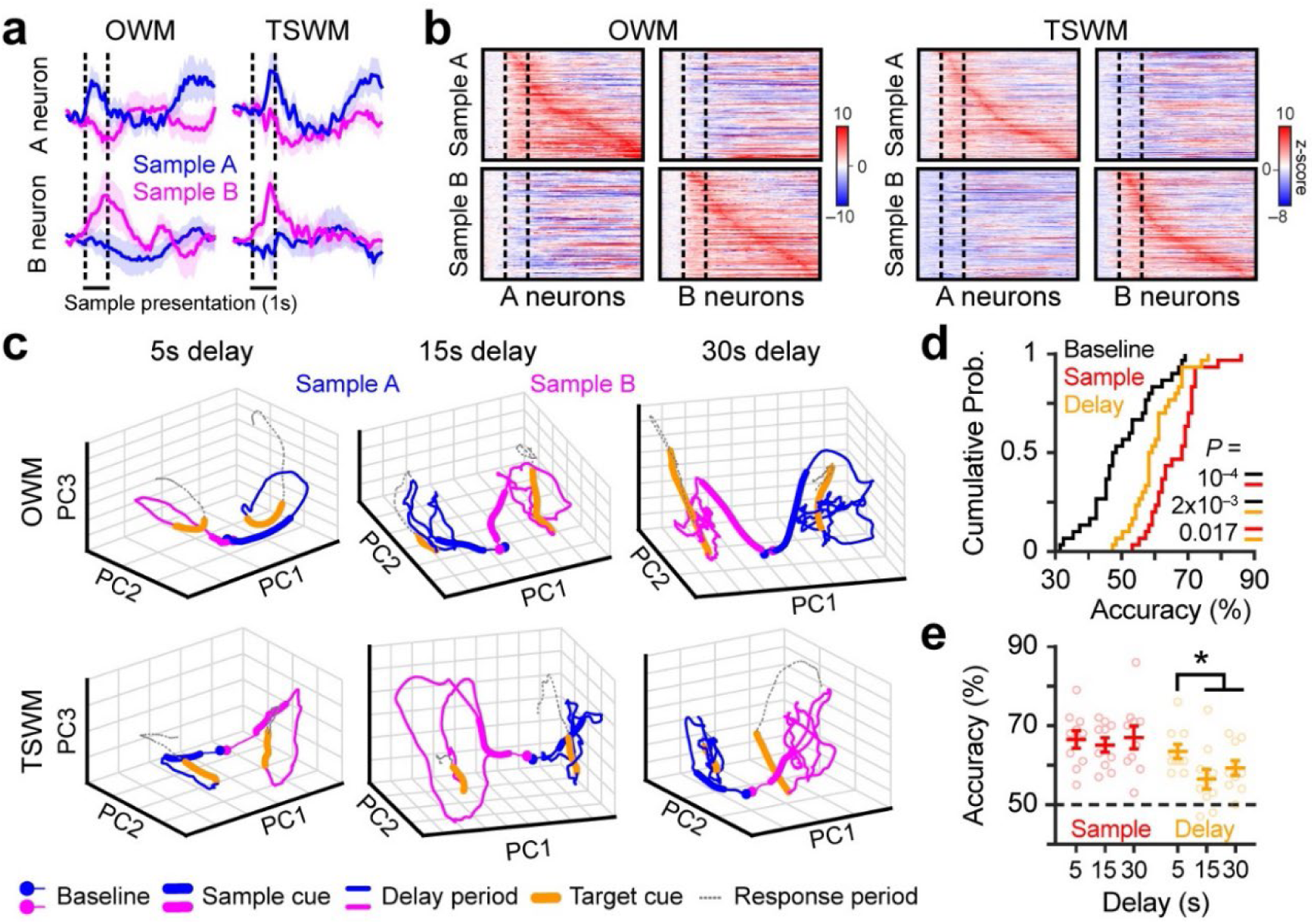
Claustrum ensembles maintain cue information during working memory. **a,** Examples of averaged calcium traces recorded upon presentation of different sample cues in four cue-selective stimulus neurons (data plotted as mean ± SD). Although each neuron exhibited an increase in calcium activity during the application of one cue but not the other, note that complex responses can still be observed during the delay period for the non-responding cue. **b,** Pseudo- color heat maps illustrating the trial-averaged responses of all cue-selective stimulus neurons. Neurons were sorted according to the latency of the peak response induced by the specific stimulus. The same neuron order was maintained for the other sample cue (*n* = 165, 150, 187, 191 neurons for samples A & B of OWM and TSWM tasks, respectively). **c,** Evolution of CLA population activity during the averaged correct trials during which either sample A or sample B was presented. Trajectories of activity are plotted in the space defined by the three principal components carrying the most variance. **d,** Cumulative probability distributions of sample cue classification accuracy evaluated at three specific time windows (baseline, sample window, and the last second of the delay; *n* = 30 mouse/delay/WM-task triplets, Kolmogorov-Smirnov test). **e,** Classification performances during the sample cue presentation and at the end of the delay period for the different delay durations (*n* = 10 mouse/WM-task pairs; repeated measures ANOVA *F*_(2,18)_ = 0.15, *P* = 0.87 and *F*_(2,18)_ = 12.4, *P* = 0.0004 for the sample and delay periods, respectively; post-hoc Fisher’s test, * means *P* < 0.05). Data are presented as mean ± SEM. See **Supplementary Table 1** for detailed statistics.

Consequently, we wondered whether CLA activity could be leveraged to match the animal’s ability to maintain cue identity in WM. For each mouse, we used the CLA population activity and a template-matching classifier to predict which of the two cues had been presented on a given trial. The accuracy of the correct prediction was independently computed per animal for each delay and task. The same classification algorithm was applied to population activity recorded during baseline, cue application, or the end of the delay. CLA activity during both cue application and the delay period significantly predicted cue identity above chance on an animal/session basis (i.e. baseline level, **Fig. 3d**). The prediction accuracy decreased after cue presentation, particularly for longer delay durations (**Fig. 3d,e**). In summary, CLA population activity retains information about cue identity, but this information degrades as the delay duration increases, thus aligning with the animals’ behavior.

### Chemo- and optogenetic inhibition of CLA neurons disrupt working memory

Can sensory information stored in CLA population activity be used for working memory functions? We explored whether selectively decreasing CLA excitability might impair WM (**Fig. 4** and **Supplementary Fig. 1**). We first expressed the inhibitory designer receptor exclusively activated by the designer drug (DREADD)^55^ hM4Di in CLA neurons of adult mice (**Fig. 4a**). Reducing the excitability of CLA neurons was sufficient to impair WM performance regardless of the sensory dimension probed (**Fig. 4b** and **Supplementary Fig. 1a**). The chemogenetic impairment induced by clozapine-N-oxide (CNO) application was transient, as WM performance returned to control levels when saline was injected the day after CNO treatment. Furthermore, the impairment was specific to DREADD activation by CNO, as control groups that did not express DREADD but received CNO did not display any WM alterations (**Fig. 4b,c** and **Supplementary Fig. 1a,b**). On average, reducing CLA excitability reduced WM performance by ∼40% irrespective of the sensory dimension tested and the transgenic mouse line used to genetically target CLA neurons (**Fig. 4c,g** and **Supplementary Fig. 1b,d**).

**Figure 4.**
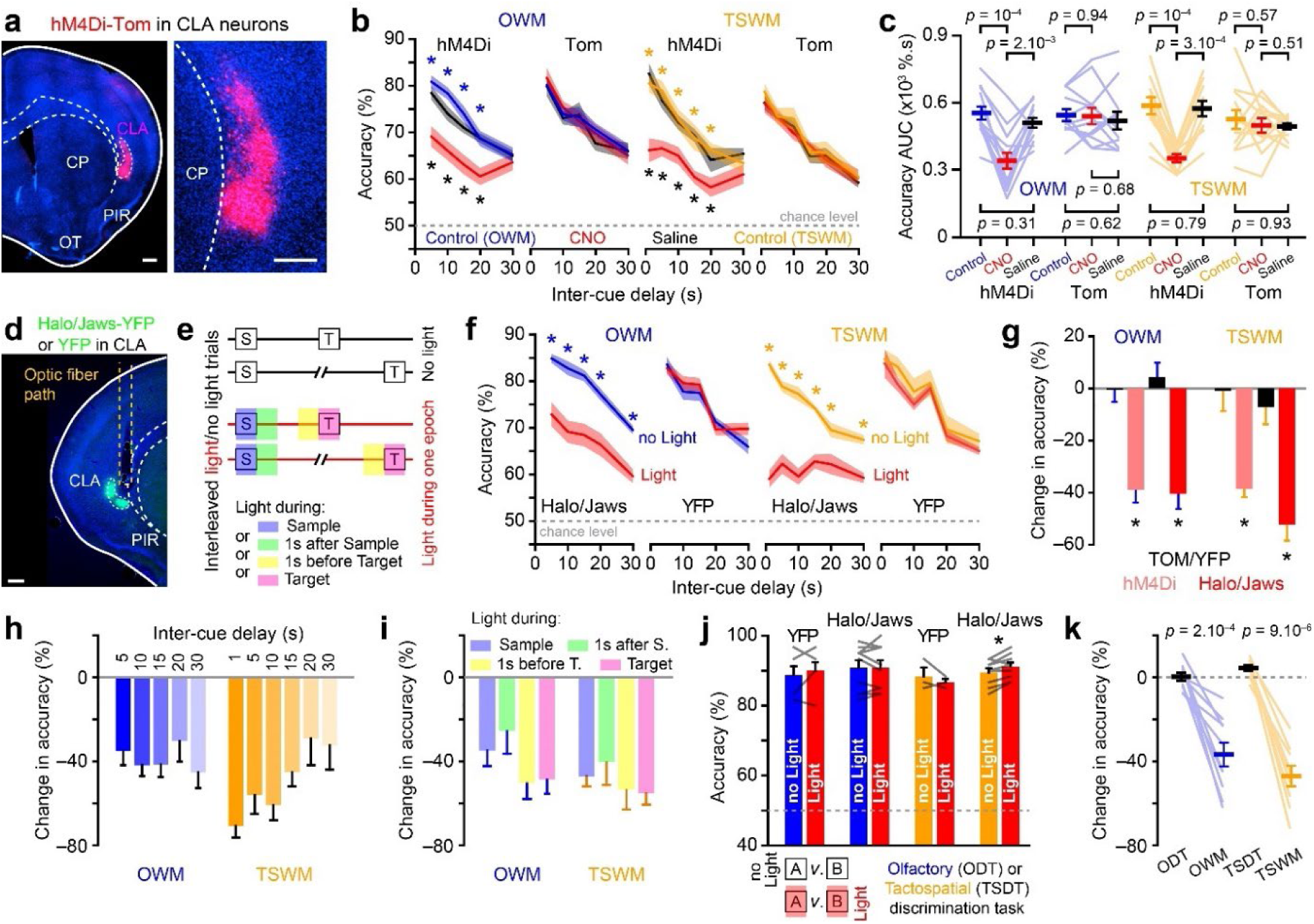
Chemogenetic and optogenetic inhibition of CLA neurons disrupt working memory. **a**, Photograph of virally targeted CLA neurons (scale bars: 250 μm). CP: caudate putamen, OT: olfactory tubercle, PIR: piriform cortex. **b,** WM performances as a function of inter-cue delay duration for mice expressing hM4Di in the CLA (OWM: *n* = 13, TSWM: *n* = 11) and control mice without hM4Di (*n* = 12 for OWM and TSWM). Control sessions (no injection), sessions with CNO injection, and sessions with saline injection were performed sequentially in this order (repeated measures ANOVA; post-hoc Fisher’s test comparisons *: *P* < 0.05, significant comparisons for Control *vs*. CNO and Saline *vs.* CNO are color-coded in blue/orange and black, respectively). **c,** Quantification of the global WM performance during the first 30 seconds measured using the Area Under the Curve (AUC) of the performance curves shown in **b**. **d,** Photograph of virally targeted CLA neurons conditionally expressing the inhibitory opsin eNpHR3.0-EYFP (scale bar: 250 μm). **e,** WM sessions with interleaved light and no-light trials across various inter-cue delays. In light trials, CLA neurons were inhibited during one of four distinct epochs (only one epoch was illuminated per behavioral session; see **Supplementary Fig. 1**). **f,** WM performance averaged across the four stimulation epochs as a function of inter-cue delay duration for mice expressing either Halo/Jaws or YFP in the CLA. Performance for light and no-light trials is compared within each group (Number of mice: Halo/Jaws [OWM: *n* = 10, TSWM: *n* = 9]; YFP [OWM: *n* = 4, TSWM: *n* = 3]; RM ANOVA; post-hoc Fisher’s test comparisons *: *P* < 0.05). **g,** Change in response accuracy (relative to control performance, i.e. no CNO or no light conditions) following either chemogenetic or optogenetic CLA inhibition in OWM and TSWM tasks shown in **b** & **f** (comparison between control and experimental groups: unpaired t-test *P < 0.05). **h,** Effect of optogenetic inhibition of CLA neurons on WM performance across different inter-cue delays (Comparison between OWM and TSWM, RM one-way ANOVA, P = 0.156). **i,** Effect of optogenetic inhibition of CLA neurons on WM performance during different light stimulation epochs for both OWM and TSWM tasks (ANOVA across light stimulation epochs, P > 0.05 for both OWM and TSWM). **j,** Performance in olfactory discrimination task (ODT) and tactospatial discrimination task (TSDT) with interleaved light and no-light trials. (Paired t-tests *: P < 0.05). **k,** Comparison of photoinhibition effects on sensory discrimination tasks (ODT and TSDT) vs. working memory tasks (OWM and TSWM) with light stimulation during the sample period (Paired t-tests; for ODT vs. OWM, P < 0.0003; for TSDT vs. TSWM, P < 0.0001). Data are presented as mean ± SEM. See **Supplementary Table 1** for detailed statistics.

While DREADD activation by CNO enables uniform inhibition across neurons expressing the engineered receptors, allowing broad modulation within the targeted population, prolonged inhibition of CLA neurons may inadvertently disrupt cortical circuit processing over extended periods, introducing potential confounding factors that complicate the determination of the specific role of the CLA in WM. Additionally, the limited temporal precision of chemogenetic approaches prevents the identification of specific time windows in which CLA activity contributes to WM (e.g. encoding *vs.* maintenance). To address this, we expressed inhibitory optogenetic actuators in CLA neurons and selectively inhibited their activity during distinct time epochs: during cue presentation, at the beginning or end of the delay period, or during target cue presentation (**Fig. 4d,e**). On average, WM performance declined in trials with light stimulation compared to interleaved no-light trials conducted in the same sessions (**Fig. 4f** and **Supplementary Fig. 1c**). Overall, WM performance was reduced by ∼40-50% in both tasks (**Fig. 4g** and **Supplementary Fig. 1d**). The impairment was modality-independent, consistent across all delay periods and specific to optogenetic inhibition, as light stimulation in the YFP control group did not negatively impact WM performance (**Fig. 4f-h** and **Supplementary Fig. 1d-f**).

To assess whether the timing of light stimulation produced distinct effects on WM performance, we analyzed each epoch individually. Inhibition of CLA neurons consistently impaired WM performance across all epochs, regardless of the specific stimulation period (**Fig. 4i** and **Supplementary Fig. 1e,g**). This effect was specific to optogenetic manipulation, as no-light trials demonstrated stable or even slightly improved performance over days, likely reflecting task mastery (**Supplementary Fig. 1e,g**). To ensure that optogenetic inhibition did not simply affect sensory processing or perception, we trained the same mice on Go/No-Go olfactory and tactospatial discrimination tasks. Comparison between light and no-light trials revealed no deficits in sensory discrimination during light stimulation (**Fig. 4j**). In contrast, performance deficits were evident during WM tasks with light stimulation at cue presentation, suggesting that the observed WM impairments were not due to sensory processing deficits (**Fig. 4k**). Overall, these findings indicate that CLA neuronal activity is crucial during cue encoding, across delay periods, and during target encoding/comparison for optimal WM performance, supporting the role of the CLA as a comprehensive WM store.

## Discussion

Our data challenges the idea that no single area contributes specifically to WM storage or that WM content is stored mainly in short-term plasticity states of synapses. If a dedicated storage area exists, then disrupting activity either during cue presentation (the encoding phase) or during any part of the delay (maintenance phase) should lead to significant information loss and behavioral impairment—precisely what we observed with optogenetic silencing of the CLA. Our findings reveal that CLA neurons exhibit persistent, content-specific activity during the delay period in WM tasks involving multiple sensory cues, where information must be retained without external inputs. This persistent activity is finely tuned to the memory content (e.g., cue identity). Disrupting CLA activity through chemogenetic or optogenetic inhibition during cue presentations or early and late delay stages consistently impaired memory retention. Notably, the same CLA manipulations in the same mice had no effect on discrimination performance, ruling out simple sensory loss due to the manipulation. Together, these data support the view that spiking activity in CLA ensembles represents working memory engrams.

Most studies concentrate on short WM delays, typically ranging from 1.5 to 5 seconds, during which sensory cortex activity often remains active and may influence information transfer. In contrast, we systematically examined the effects of chemogenetic and optogenetic manipulations across delays ranging from 1 to 30 seconds, thereby enhancing our understanding of WM maintenance over longer durations. Regardless of the sensory dimension tested, reducing CLA excitability resulted in a significant decline in WM performance, averaging approximately 40-50%. In some cases, performance dropped by as much as 70% for specific delays, with a complete shutdown observed in some individual mice. The impact of these manipulations was notably strong, particularly given two key factors: first, our interventions decreased excitability without completely suppressing firing activity across the entire neuronal population; second, our estimates indicate that we affected at most 50% of the entire CLA population due to limitations in viral infection that hindered full anteroposterior coverage^8,9^ and constraints of the Cre lines in targeting the entire population of CLA projection neurons^56^. It is essential to emphasize that while we propose the CLA as a critical storage site for WM, we do not claim it to be the sole region capable of maintaining some WM information. Consequently, the residual WM performance observed during chemogenetic and optogenetic treatments may suggest the involvement of additional, albeit limited, storage sites within the brain.

By integrating results from imaging, chemogenetic, and optogenetic interventions, we propose that early-stage activity induced by cue presentation drives persistent attractor-like states within the CLA. Disrupting this persistent state at any point during WM tasks irreversibly alters the neural dynamics essential for maintaining WM. The consistent effects observed across all delays and light stimulation epochs also rule out the possibility of neural redundancy, where the inhibition of one area could be compensated for by other regions or pathways involved in the task. Our study provides compelling evidence that the CLA plays a critical role in WM by sustaining the persistent activity necessary for task performance. The application of temporally precise optogenetic inhibition underscores the importance of CLA activity throughout different phases of the WM task, highlighting its essential function in maintaining working memory. These findings enhance our understanding of the neural mechanisms underlying WM and suggest that the CLA serves as a pivotal local storage buffer, unifying sensory information to guide behavior. However, the mechanisms underlying cue-specific responses and their persistence in CLA neurons remain to be fully elucidated. It is likely that the cortex drives CLA activity during cue presentation and part of the delay period, with a portion of the sustained activity arising from intra-CLA connectivity. While initial theories suggested local interactions among CLA projection neurons primarily occur through feedforward inhibition^57^, recent data reveal robust glutamatergic connectivity between these neurons^58^. Future studies should explore the precise pathways responsible for conveying specific information to the CLA and the mechanisms governing activity maintenance.

In conclusion, we propose an analogy in which the CLA functions as a volatile memory store, akin to a computer’s Random Access Memory (RAM), serving as short-term, rapidly accessible storage. The primary role of RAM is to provide fast, temporary storage for data actively in use, allowing seamless task processing. In this analogy, different cortical areas act as parallel Central Processing Units (CPUs) that generate and load information into the RAM (CLA), enabling swift retrieval and manipulation as needed. Just as a computer’s performance improves with more RAM, allowing for the simultaneous storage of larger amounts of information without slowdown, this model could explain the limited capacity of items stored in working memory, which would be constrained by the CLA’s storage capacity. When the CLA’s storage would be full or subject to interference from competing sources of information, working memory performance would decline.

## Acknowledgments

We thank members of A.C. and I.R. laboratories for helpful discussions and/or comments on the manuscript. We thank Sébastien Pellat, Elodie Husi, Chenda Kan, Véronique Jungo, and Loris Mannino for expert technical assistance. This research was supported by the University of Geneva and the European Research Council (contract ERC-SyG-856439-CLAUSTROFUNCT to A.C. and I.R.), the Swiss National Science Foundation (grant 310030_215572 to A.C.) and the Novartis foundation for medical research (A.C. and I.R).

## Author contributions

A.C., A.S.B., and I.R. conceptualized and designed the study. A.S.B. conducted viral injections, behavioral testing, calcium imaging experiments, and all subsequent analyses. C.H. assisted with calcium imaging and analysis. B.U. and J.-R.R. provided support for the behavioral experiments. A.C., A.S.B., and I.R. collaborated on writing and editing the manuscript, incorporating feedback from all authors.

## Declaration of interests

The authors declare no competing financial interests.

## Methods

### Animals

A total of 44 mice (22 males and 21 females), aged between 3 and 11 months (at then of some experiments), were included in the study. All experiments were conducted using either *Slc17a6^tm^*^2^*^(cre)Lowl^*/J heterozygote mice (referred to as *Vglut2*-cre in the text; the Jackson Laboratory, strain number 016963), Tg(*Smim32*-cre)61*^Irod^* heterozygote mice (referred to as *Smim32*-cre in the text; 5 males and 9 females; a detailed characterization of this line is available at ^56^), or Tg(*Smim32*-cre)61*^Irod^*;*Gt(ROSA)26Sor^tm^*^1^*^(CAG-CHRM^*^4^*^*,-mCitrine)Ute^*/J double heterozygote mice, which were obtained by crossing *Gt(ROSA)26Sor^tm^*^1^*^(CAG-CHRM^*^4^*^*,-mCitrine)Ute^*/J homozygote mice^59^ (the Jackson Laboratory, strain number 026219) with Tg(*Smim32*- cre)61*^Irod^*. In *Vglut2*-cre mice, the cre coding sequence does not disrupt the *Slc17a6* gene coding sequence since it was placed after the stop codon^60^. Housing conditions involved maintaining mice in groups of 2-5 individuals, following a standard 12h light/dark cycle, within a room maintained at 24°C, with *ad libitum* access to food. Prior to behavioral experiments, mice underwent a 12-14h period of water restriction. After the behavioral sessions, mice were provided unrestricted access to water until satiety within their housing cages. All experiments were conducted during the light phase of the light/dark cycle. All experiments adhered to the ethical guidelines outlined in the Swiss Federal Act on Animal Protection and the Swiss Animal Protection Ordinance. The experimental protocols received approval from the University of Geneva and the Geneva state ethics committees (Licenses GE/94/20 and GE368).

### Surgical procedures and animal preparation

#### Preparation of mice for surgeries and post-operative care procedures

Mice were anesthetized using isoflurane at concentrations of 3-5% for induction and 1.5-2% for maintenance. Prior to any intervention, the skin overlying the skull was disinfected with a betadine solution and subsequently removed under local anesthesia using 0.25% carbostesin. To mitigate the risk of cortical edema during craniotomy, mice were administered an intraperitoneal injection of dexamethasone (4 mg/ml). For the procedure, mice were securely positioned using ear bars and a nose clamp on a stereotaxic apparatus (Stoelting) to ensure stability and precision. To prevent ocular dehydration, artificial tears (Lacryvisc™ eye gel) were applied, while the body temperature was monitored and maintained at ∼37°C using a heating pad. Specific surgical procedures were then performed as dictated by experimental requirements (see details below). Following completion, to counter any dehydration during anesthesia, a subcutaneous injection of 0.9jM NaCl (∼500 µl) was administered. Furthermore, post-operative care included the provision of analgesia through a subcutaneous injection of carprofen (1 mg/ml) and the addition of paracetamol (2 mg/ml) to the drinking water for three days.

#### Adeno-associated virus injections

AAV2-hSyn-DIO-hm4D(Gi)-mCherry (Titer GC/ml: 4.6x10^12^, Addgene catalog #: 44362-AAV2), AAV1-Syn-Flex-GCaMP6f.WPRE.SV40 (Titer GC/ml: 1.9x10^13^, Addgene catalog #: 100833-AAV1), AAV1-syn-FLEX-jGCaMP8f-WPRE (Titer GC/ml: 5.9x10^12^, Addgene catalog #: 162379-AAV1), AAV2-Ef1a-DIO-EYFP (titer: 4.5 × 10¹² GC/ml; UNC GTC vector core; catalog #: AV4842F), AAV1-Ef1a-DIO-eNpHR3.0-EYFP (titer: 2.1 × 10¹³ GC/ml; Addgene catalog #: 26966-AAV1), and AAV1-CAG-FLEX-rc[Jaws-KGC-GFP-ER2] (titer: 2.5 × 10¹³ GC/ml; Addgene catalog #: 84445-AAV1) were used in the experiments. Following the preparation of mice for surgical procedures, craniotomies were performed either unilaterally or bilaterally above the CLA using an air- pressurized driller. Glass pipettes filled with AAV solutions were inserted in one hemisphere at a time at the following coordinates: antero-posterior (AP) 1.1 mm and medio-lateral (ML) ± 2.85 mm relative to bregma, dorso-ventral (DV) -2.4 mm relative to the brain surface. Approximately 100 nL of the AAV solution was deposited per injection site. The pipette remained in place for 5 minutes before removal. For chemogenetic and optogenetic experiments, infections were bilateral, while for microendoscopic imaging experiments, infections were unilateral. Following viral injections, additional surgical procedures were performed based on the specific experiment requirements. A minimum period of three weeks was allowed for viral expression before mice underwent behavioral experiments.

#### Headpost implantations

After securing the mouse on the stereotaxic apparatus, the overlying skin was removed, and the surgical site was cleaned with sterile saline. The periosteum and exposed muscles were gently scraped off from the skull, which was then allowed to air dry. Gelfoam® soaked in sterile saline was applied in case of minor bleeding to facilitate clotting. The skull was chemically etched using a dental etching agent to enhance surface roughness for better adherence to dental cement. Any excess etching agent was cleaned with saline, and the surface was air-dried. A thin coat of cyanoacrylate gum (Cyberbond) was applied to prevent fluid seepage and reinforce the skull for improved cement adherence. A dental primer was applied, air-dried, and polymerized with UV light. UV-polymerizing cement was then spread over the skull, fused with the periphery and polymerized. The head-post, coated with UV-polymerizing cement, was positioned and polymerized. Dental acrylic cement, mixed with a polymerizing agent and cyanoacrylate gum, was used to seal exposed areas on the skull.

#### GRIN lens implantation

After preparing the mouse for surgery, a craniotomy was performed over the CLA. Following a unilateral injection of AAV1-Syn-Flex-GCaMP6f.WPRE.SV40, a 0.5 mm diameter GRIN snap-in imaging cannula (Model L, Doric Lenses Inc., Canada) was gradually lowered into the brain by approximately 2.45-2.55 mm. The GRIN cannula was securely affixed to the skull using cyanoacrylic glue (Vetbond™) and dental cement. Following the GRIN lens assembly, a head-post was attached to the skull following the aforementioned procedures. Mice were allowed a recovery period of at least three weeks post-surgery.

### Delayed Non-Match to Sample (DNMS) working memory tasks

#### General description

Prior to starting behavioral experiments, the mice underwent a brief water restriction period lasting 1-2 days. Before each session, mice were gently placed in a plastic/PVC tube and their head was securely restrained using a specialized metallic device affixed to a stable platform, as previously described^61,62^. All experiments were conducted using custom-made, computer-controlled olfactometers, following established protocols^63,64^. These olfactometers were designed to ensure precise delivery of sensory stimuli (tested with photoionization detectors ^65^), including odorants or a clean airflow, and to accurately measure lick responses within predetermined time windows^61^. This was achieved through the integration of a licking circuit and a metallic water port.

In the DNMS WM tasks, each trial began with a 2s baseline, followed by the presentation of a sample cue for 1s, succeeded by a variable delay period, and then a target cue presented for 1s, which could either be the same (match trial) or differ from the sample cue (non-match trial). Mice underwent training to initiate licking responses specifically within a response window during non-match trials (see next section). The 2.5s response window started 0.5s following the onset of the target cue delivery. Trials were separated by a fixed inter-trial interval of 10s. To help the mice distinguish between the inter-trial cue presentation and the intercue delay period, a 0.2s tone, with a frequency ranging between 3 and 9 kHz depending on the animal, was used to indicate the sample cue for each trial. As spontaneous licking occasionally occurred, licking events were recorded throughout the entire trial duration. In instances where mice exhibited licking prior to the designated response window, a higher count of licks during the response window was required in non-match trials to qualify as a ’Hit’ (correct) trial. Upon meeting the licking criterion successfully, mice were promptly rewarded with water delivery (approximately 5 µl). A ’False Alarm’ occurred when licking events were detected in the response window during match trials. Importantly, mice did not face any punishment in False Alarm trials. Neither rewards nor punishments were administered for ’Miss’ (no-lick in a non-match trial) or ’Correct Rejection’ (no-lick in a match trial) trials. Each day, mice completed approximately 100 trials, with all these trials being utilized for computing behavioral results. Trial-related events, such as stimulus onset and offset, licking events, and water delivery, were systematically recorded by computers.

In this study, we assessed WM across distinct sensory modalities, namely, olfactory WM (OWM)^18,66,67^ and tactospatial WM (TSWM). As our focus was on OWM rather than olfactory perception or discrimination, we used various monomolecular odorants (ethyl acetate, 2-pentanone, 1,4-cineole, eugenol, amyl acetate and ethyl butyrate; Sigma-Aldrich) known to be easily discriminated by mice^48,61,68^ and not requiring a functional reorganization of olfactory responses such as pattern separation associated to learning^69,70^. The odorants were initially diluted in mineral oil (2% odorant dilution) and further mixed with a clean carrier airflow (odorant: carrier ratio of 1:2), ensuring activation of multiple olfactory bulb glomeruli^68,71^ without eliciting the large responses observed following natural odorant application^72^. We did not synchronize odor application with respiration to minimize equipment crowding near the mouse snout. Consequently, we opted for a 1s odor application to ensure at least two complete respiratory cycles during odorant presentation while minimizing strong olfactory afterimages in the olfactory system, which could complicate WM performance comparisons across sensory modalities^65^. For the same reason, we did not test the 1s inter-cue delay in the OWM task to reduce the potential impact of residual olfactory afterimages as confounding factors in WM performance.

Counterbalancing was employed to evenly distribute the selected odorants as sample and target cues across different trials. In the TSWM task, we used identical equipment to administer a gentle air puff (400 ml/min) directed towards either the left or right whisker pad (**Fig. 1a**). During each trial, if the air puff was directed to the same whisker pad for both the sample and target cues, it was categorized as a match trial. Conversely, if the air puff was delivered to opposite whisker pads, the trial was classified as non-match. Additionally, we implemented counterbalancing of the air puff direction as sample and target cues across different trials.

#### Behavioral training

The comprehensive behavioral experimental process included habituation, pre-training, training, and the evaluation of working memory through DNMS tasks. During habituation, mice were gently head-fixed in behavioral setups and introduced to drinking water from the lick port, aided by manually delivered water. Once mice became accustomed to head restriction, the pre-training began. In the initial phase (20 trials), a water drop (2-3 µl) was promptly presented to mice following a 0.2s tone. Subsequent trials introduced increasing delays between the tone and water reward, progressing to 2s (10 trials) and 5s (10 trials), aimed at eliciting licking behavior and dissociating the tone from water delivery. Upon mastering the ability to wait for water rewards, pre-training shifted focus exclusively to non-match trials. Ethyl acetate and 2-pentanone served as stimuli, randomly assigned as sample and target cues across trials. Sample and target cues were presented with a fixed intercue delay of 5s. In the initial 20 trials of this phase, an automatic reward was delivered at the onset of the response window. Following this, the automatic reward was withheld, and any licking event during the response period triggered water delivery (20-40 trials). Once mice started licking regularly in the response period, the task rules were implemented. A correct trial now necessitated a higher count of licks during the response window compared to the pre-response window. Typically, mice took approximately 150-200 trials to learn the task rules and consistently exhibited licking behavior during the response window. Following this, the training phase for OWM was initiated, introducing a pseudorandomized implementation of both match and non-match trials. During this phase, 1,4-cineole and eugenol were used as cues and the intercue delay was maintained at 5s. Specifically, one or two non-match trials and one or two match trials were presented in a randomly sequenced pattern, appearing in blocks of two or four trials. Subsequently, the trials were pooled together, and performance accuracy was determined by calculating the percentage of correct trials within each block of 100 trials. The performance accuracy, denoted as ’accuracy (%)’ in figure labels, is defined as follows:

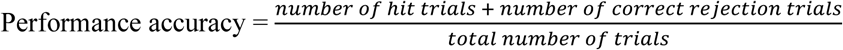

In addition to performance accuracy, we extended our evaluation to include the Discriminability Index (D-prime). This index quantifies the ability to discern the relevant signal from background noise, offering a quantitative measure of mice’s capability to make accurate discriminations. The calculation of D-prime involved determining the Hit Rate (HR) and False Alarm Rate (FA) as follows:

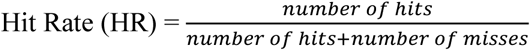

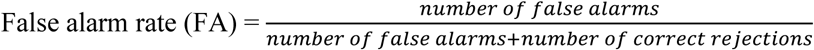

Discriminability is then defined by the formula:

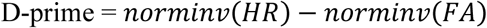

where *norminv* is the inverse of the cumulative normal distribution function. To prevent issues related to infinite values, we applied a conversion to Hit and False Alarm rates. If either rate equaled 100%, it was adjusted to 0.99, and if the rate was 0, it was set to 0.01 during the conversion process.

In most instances, mice learned to perform the task within 200 to 300 trials (**Fig. 1b**). To ensure the stability of their acquired skills, training was extended up to 800 trials. Once mice were proficiently trained, they were exposed to DNMS tasks with variable delay periods, all within a single session. The delay periods were pseudo-randomly varied in blocks of 2 or 4 trials, encompassing delay periods of 5s, 10s, 15s, 20s, 30s, 40s, and 60s. Each session consisted of 100 trials, with each delay period presented for 14-16 trials within the session. By maintaining a fixed number of trials at 100, we ensured that each mouse completed all trials within a single session. To bolster the robustness of our observations, mice underwent three such sessions across three different days. This approach not only facilitated a comprehensive investigation of the mice’s response to variable delays but also enabled a thorough analysis by ensuring an adequate number of trials for each specific delay condition. In order to assess the generalizability of performance and ensure that results were not solely odor-specific, 12 out of the total 30 mice additionally underwent testing with amyl acetate and ethyl butyrate. Following confirmation of consistent performance in the DNMS task across various odors, performance was averaged across different odor sessions. This averaged performance data was then utilized for the analyses presented in **Figure 1**.

In the TSWM DNMS tasks, mice were trained with a 1-second delay between sample and target stimuli. As TSWM training followed the completion of OWM tasks, mice had already mastered the procedural elements of the task paradigm. TSWM training specifically aimed to familiarize mice with the shift in sensory modality, teaching them to detect and distinguish the directionality of whisker pad stimulation. Remarkably, all mice successfully adapted to this sensory shift within the initial 100-150 trials. To ensure proficiency in TSWM, all mice underwent training for up to 600 trials. Additionally, for TSWM with variable delays, an extra delay of 1 second was introduced alongside the delays used for OWM. All other aspects of the task design remained consistent with those of the OWM.

The accuracy and D-prime values of the WM tasks at varying delay durations were modeled using an exponential decay function in GraphPad Prism 9.0. We fixed the plateau to a constant value, set at 50 for accuracy and 0 for D-prime values, representing the theoretical chance level for the behavior. The function was defined as follows:

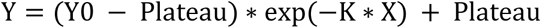

Using this equation, we calculated Y0, representing the extrapolated amplitude at X = 0, and the half-life, computed as ln(2)/*K*.

### Microendoscopic calcium imaging

#### Image acquisition and image processing

Imaging acquisition utilized a snap-in fluorescence microscope body (2CFM model L Type D, Doric Lenses Inc., Canada) mounted on the implanted GRIN cannula. The Ce:YAG fiber light source (465 nm output, Doric Lenses Inc., Canada) was adjusted within the range of 1-2.7 mW. The field of view, corresponding to 320 µm x 320 µm, was captured at a spatial resolution of 600 pixels x 600 pixels and a frame rate of 10 Hz. Image processing involved the use of ImageJ (National Institutes of Health, USA), Doric Neuroscience Studio (Doric Lenses Inc., Canada), and custom MATLAB (MathWorks, Inc.) routines. On imaging days, mice completed the entire session of 100 trials, with only the 5s, 15s, and 30s delay trials being imaged. Synchronization with the trials was achieved through an external TTL signal from the olfactometer software. Images were acquired over three days to ensure an adequate number of trials for each specific delay condition.

Images from different trials of a given session were concatenated using a custom script written in MATLAB. Individual image frames were corrected for infocal (XY) plane brain motion using the alignment function integrated into Doric Neuroscience Studio and the maximum projection of the concatenated trace served as a reference for the motion correction. Following motion correction, the edges of the images were cropped by 5 pixels, and the trimmed images were downsampled by a factor of two using bilinear interpolation in MATLAB. Additionally, a Kalman stack filter in ImageJ was applied to remove high-gain noise from the images. To identify neurons and detect Ca^2+^ transients, we used a cell-sorting algorithm^7^ implemented in MATLAB. This algorithm integrates principal and independent component analyses (PCA-ICA) and is coupled with a segmentation routine optimized to minimize the occurrence of false positives. To remove slow trends and preserve faster dynamics in the signal, we applied discrete wavelet transformation to the traces from each neuron. After detrending, the traces were split into individual trials. For normalization, a baseline vector was constructed from all trials of a given neuron, with the baseline defined as the period 1s before sample presentation. The median of this baseline vector was then used to calculate z-scored traces as follows:

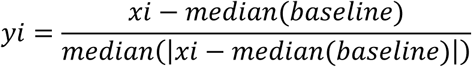

Neurons recorded from different mice were grouped to generate a high-dimensional neuronal population. The classification of neurons into distinct categories, namely Stimulus neurons, Delay neurons, and Inhibited neurons, was performed using the averaged activity derived from correct trials for all neurons. This categorization relied on the time at which the response amplitude first surpassed a predefined threshold of one. Specifically, if the threshold was first crossed during the sample presentation, the neuron was classified as a Stimulus neuron. Alternatively, if the threshold was surpassed after the offset of the sample presentation, the neuron was designated as a Delay neuron. Neurons that did not cross the threshold were categorized as Inhibited neurons. This classification scheme offers a nuanced understanding of the temporal dynamics and response characteristics exhibited by different neuron populations (**Fig. 2b,d**). The activity heat maps were generated by plotting the Inhibited neurons first, followed by the Stimulus neurons, and then the Delay neurons and visualized as a pseudo-color heat map using MATLAB (**Fig. 2c**).

We did not attempt to compare the representations evoked by cues across different sensory modalities due to the separation of imaging sessions by several weeks, raising uncertainty about whether the sampled CLA neurons were consistent across these time points.

To gain insights into the trial-dependent population response dynamics, temporal correlations were calculated for four distinct trial structures (sample-target): A-A, A-B, B-A, and B-B. To compute the temporal correlation matrices, a population vector with N dimensions was constructed, where N denotes the total number of responsive neurons recorded across all mice. Within each dimension, we calculated the mean calcium z-score amplitude averaged across all correct trials for all task-delay pairs every 100 ms. Consequently, the temporal evolution of CLA population activity was generated by calculating pairwise Pearson’s linear correlations between each pair of time bins, where each time bin corresponds to a duration of 100 ms. The resulting correlation matrix was visualized as a pseudo-colored heat map (**Fig. 2f**). To further grasp the temporal progression of population activity, we compared all time vectors to those recorded at the peak response time of stimulus neurons and at the conclusion of each delay period (**Fig. 2g,h**).

A two-step analysis was undertaken to identify neurons with stimulus-specific responses (i.e., sample A or sample B in **Fig. 3a,b**). Firstly, the average responses were calculated from the correct trials where either stimulus A or stimulus B served as the sample for each neuron separately. Subsequently, for each neuron, a two-sided Wilcoxon rank sum test was conducted, comparing the activity during the sample presentation to discern whether the neuron exhibited specificity to stimulus A or B. The neurons displaying specificity to a particular stimulus were then visually represented through a pseudo-colored heat map. These selective neurons were sorted based on the latency of their maximum activity to enhance interpretability. This particular ordering of neurons was maintained to visualize their responses toward the non-specific stimulus. For the analysis of population trajectories, the averaged responses of all neurons toward stimuli A and B (from correct trials) were concatenated to construct a data matrix with rows representing individual neurons and columns representing time points. To visualize this high-dimensional data, we performed Principal Component Analysis (PCA), and the resulting first three Principal Components (PCs), which are eigenvectors with the largest eigenvalues of the covariance matrix of the original firing patterns, were used to generate trajectories shown in Figure 3c.

To assess classification performance for the sample stimulus, a meticulous process was employed, involving the following steps. This classification analysis was conducted on individual mice with a minimum of 10 recorded neurons, ensuring a substantial dataset for accurate classification assessments (n = 5). For each mouse, responses were averaged across neurons, generating a single response profile for every trial. Specifically, correct trials where stimulus A served as the sample (either A-A or A-B) were aggregated to create the template response A, and the same process was followed to create template response B. These templates were derived from different temporal windows, including the baseline (1 second before sample presentation), sample period (1 second duration), or the end of the delay period (1 second before target presentation). To execute the classification, one trial was selected as the test trial. If this trial contributed to the creation of either template response, the template was recalculated by excluding the test trial, thereby preventing oversampling bias in the classification process. Pairwise Pearson’s Linear Correlation Coefficient between the test trial (extracted from the same temporal window as the template response) and the response templates was then computed. Subsequently, trials were assigned to the closest template, effectively predicting the stimulus category.

### Chemogenetic manipulation

We employed two distinct approaches to silence neuronal activity during behavior. In the first group of mice, we injected bilaterally into the CLA of *Vglut2*-cre mice AAVs that conditionally express Designer Receptors Exclusively Activated by Designer Drugs (DREADDs AAV2- hSyn-DIO-hM4Di-mCherry). These viruses were the Gq-coupled DREADDs that employ a modified form of the human M4 muscarinic receptor (hM4Di) to induce inhibitory cellular responses in the presence of the ligand clozapine-N-oxide (CNO)^8^. This method allowed us to achieve a strong inhibition of neurons around the infected site. However, considering the elongated structure of the CLA, the second approach was adopted. *Smim32* is a specific marker of CLA neurons in mice (unpublished single-cell RNA sequencing data from our laboratories) and in humans^73^. Thus, we employed Tg(*Smim32*- cre)61*^Irod^*;*Gt(ROSA)26Sor^tm^*^1^*^(CAG-CHRM^*^4^*^*,-mCitrine)Ute^*/J heterozygous mice to enable the expression of the inhibitory DREADD in approximately 50% of the entire CLA population. This alternative approach provided a more widespread modulation of neuronal activity across the CLA, offering a complementary perspective to the localized inhibition achieved in the first approach.

In the chemogenetic inhibition experiments, mice participated in a sequence of OWM and TSWM sessions. The experimental timeline involved three initial sessions without treatment, establishing a baseline for the mice’s behavioral performance. Subsequently, mice received CNO treatment for three sessions, administered over three consecutive days. Following the CNO treatment, an additional three sessions were conducted with saline administration. The inclusion of saline sessions served as an internal control, enabling the tracking of behavior both before and after activating the DREADD receptors. This experimental design allowed us to examine the impact of chemogenetic inhibition on WM across distinct sensory modalities. The chemogenetic manipulation was carried out 30 minutes after the intraperitoneal injection of CNO at a dosage of 2 mg/kg (TOCRIS Bioscience) or after the administration of saline. By incorporating both treatment and control sessions, we could discern any alterations in cognitive performance attributed to the specific modulation of neuronal activity induced by the chemogenetic intervention. Moreover, to control for any potential non-specific effects of CNO alone on WM, we performed parallel experiments with *Vglut2*-ires-cre mice injected unilaterally in the CLA with AAV1-Syn-Flex-GCaMP6f.WPRE.SV40 or pGP-AAV1-syn-FLEX-jGCaMP8f-WPRE. These mice, lacking DREADD expression, served as a DREADD-free control group. They underwent the same training regimen, including initial sessions without any treatment, followed by CNO and saline sessions. To assess the impact of CNO on WM, we quantified the Area Under Curve (AUC) for the WM curves. Specifically, the AUC measurement spanned from 5s to 30s for OWM tasks and from 1s to 30s for TSWM tasks. The AUC estimation was carried out using GraphPad Prism 9.0.

For the OWM, the *hM4Di* group was composed of 7 *Vglut2*-cre mice and 6 *Smim32*-cre mice expressing hM4Di; 9 males and 4 females), in the *no hM4Di* group (11 *Vglut2*-cre mice expressing GCaMP indicator; 8 males and 3 females, and 1 female *Smim32*-cre negative), repeated measures ANOVA F_(2,30)_ = 0.102, P = 0.904. For TSWM, in the *hM4Di* group (6 *Vglut2*-cre mice and 5 *Smim32*-cre mice expressing hM4Di; 8 males and 3 females), in the *no hM4Di* group (11 *Vglut2*-cre mice expressing GCaMP indicator; 8 males and 3 females, and 1 female *Smim32*-cre negative), repeated measures ANOVA F_(2,33)_ = 0.355, P = 0.705.

### Optogenetic manipulation

#### Viral infections and optical fiber implantation

*Smim32*-cre mice were divided into two groups for the optogenetic experiments. The control group (YFP) included four mice injected with AAV2-Ef1a-DIO-EYFP. The experimental group (Halo/Jaws) consisted of animals injected with either AAV1-Ef1a-DIO- eNpHR3.0-EYFP or AAV1-CAG-FLEX-rc[Jaws-KGC-GFP-ER2] (5 mice each). The preparation of mice for surgical procedures, virus injections, and headpost implantations is mentioned in the above section. Following viral injections, two optical fibers (200 μm diameter, 0.39 NA) with ceramic ferrules were implanted bilaterally, positioned 300 μm above the virus injection sites. Optical fibers with a transmission efficiency greater than 70–80% were selected for implantation. A mixture of dental acrylic cement and cyanoacrylate glue was applied around the ceramic ferrules to provide structural support.

#### Laser illumination design in optogenetic experiments

Viral expression was allowed to proceed for at least four weeks before initiating laser illumination. A bifurcated optical fiber bundle (200 µm core, 0.39 NA) was coupled to the implanted optical fibers using ceramic sleeves. Laser power at the end of the external fiber was measured with a laser power meter (Thor Labs, PM100D) and adjusted to achieve maximum illumination without raising the local tissue temperature^74^Click or tap here to enter text.. Illumination was provided with Ce:YAG fiber light source (Doric Lenses Inc., Canada). For YFP and Halo groups, a 559 nm or 563 nm bandpass filter was used to produce yellow light, while a 612 nm bandpass filter produced orange light for the Jaws group. Illumination was controlled using Doric Neuroscience Studio (Doric Lenses Inc., Canada). Laser power was set to 5-6 mW for the Halo and YFP groups and to 12 mW for the Jaws group.

In optogenetic inhibition experiments, mice participated in a series of OWM and TSWM sessions with different light-stimulation protocols. Each 100-trial session included an equal number of light-stimulation trials (light) and non-stimulation trials (no-light), arranged in interleaved pseudorandomized blocks of 2–4 trials. In light trials, CLA neurons were inhibited during one of four specific time epochs: (1) during the Sample presentation, (2) for 1s after the Sample presentation, (3) for 1s before the Target presentation, or (4) during the Target presentation (**Fig. 4e**). Only one of these time epochs was targeted per daily session, and each epoch was repeated across six consecutive sessions before moving to the next. This design ensured sufficient trial counts across all delays for both light and no-light conditions. An exception to this protocol was made for post-Sample light stimulation in the OWM task (first design tested), where light and no-light trials were conducted in separate sessions (three sessions each). This slight adjustment in stimulation strategy did not affect the overall accuracy difference between light and no-light trials (**Fig. 4i**). To control for visual interference, a masking light of the same wavelength as the stimulation laser was flashed continuously above the animals’ head for both light and no-light trials. The effect of photoinhibition on WM performance was calculated as the percentage change in accuracy within a session, using the following formula:

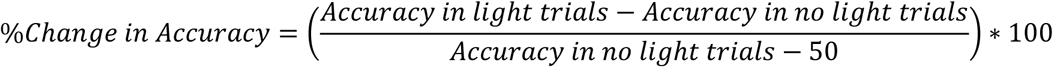

To evaluate the effect of photoinhibition on sensory discrimination, a Go/No-Go task was designed for both olfactory (1,4-cineole vs. eugenol) and tactospatial (gentle air puff directed toward either the left or right whisker pad) modalities. Each trial began with a tone, followed by the 1-second sensory cue. The response period coincided with cue presentation and continued for 2 seconds, with a 10-second inter-trial interval. Mice were trained to lick in response to the Go stimulus. In Go trials, a lick during the response period was recorded as a Hit, while no lick was marked as a Miss. In No-Go trials, a lick during the response period was counted as a False Alarm, while no lick was a Correct Rejection. Hits were rewarded immediately with 2–3 µl of water, with no rewards or penalties for other responses. Go and No-Go trials were pseudorandomized, allowing no more than two consecutive trials of the same type. Training continued until mice reached performance levels above 80%. Once trained, mice completed a 120-trial session with interleaved light and no-light trials, balanced evenly across both conditions.

### Histology

To validate the viral infections, lens or optic fiber implantations, mice underwent transcardial perfusion with 25 ml of phosphate-buffered saline (PBS) followed by 25 ml of 4% paraformaldehyde (PFA) in PBS. Subsequently, brains were dissected, post-fixed in 4% PFA for 24h, and then stored in PBS. To visualize the expression patterns, consecutive 50 µm coronal sections were sliced using a vibratome (Leica VT1000S). The tissue sections underwent a washing regimen involving three 15-minute cycles with PBS. Staining with Hoechst (1:5000, Invitrogen, cat no: H3570) was performed for 15 minutes to highlight cell nuclei, followed by another 15-minute wash in PBS. Slices were mounted on slides, embedded in a mounting medium (Vectashield, Reactolab S.A.), and recovered by a thin glass coverslip. The samples were imaged using a ZEISS Axiocam multi-channel fluorescence imaging microscope using either a 4X or 10X objective. Typically, the expressed fluorophore exhibited the distinctive ’peanut’ shape of the CLA, as illustrated in **Fig. 4a,d**.

### Statistics

Statistical analyses were performed using MATLAB or GraphPad Prism 9.0. The study utilized ANOVA in conjunction with post-hoc multiple comparison tests, the Kolmogorov–Smirnov test, and the Wilcoxon signed-rank test. All tests were two-sided, and unless otherwise specified, error bars in the figures represent the standard error of the mean.

#### Reporting Summary

## Code availability

The custom-written MATLAB scripts will be made public on a repository.

## Data availability

The data that support the findings of this study will be made public on a repository.

---

**Supplementary Figure 1.**
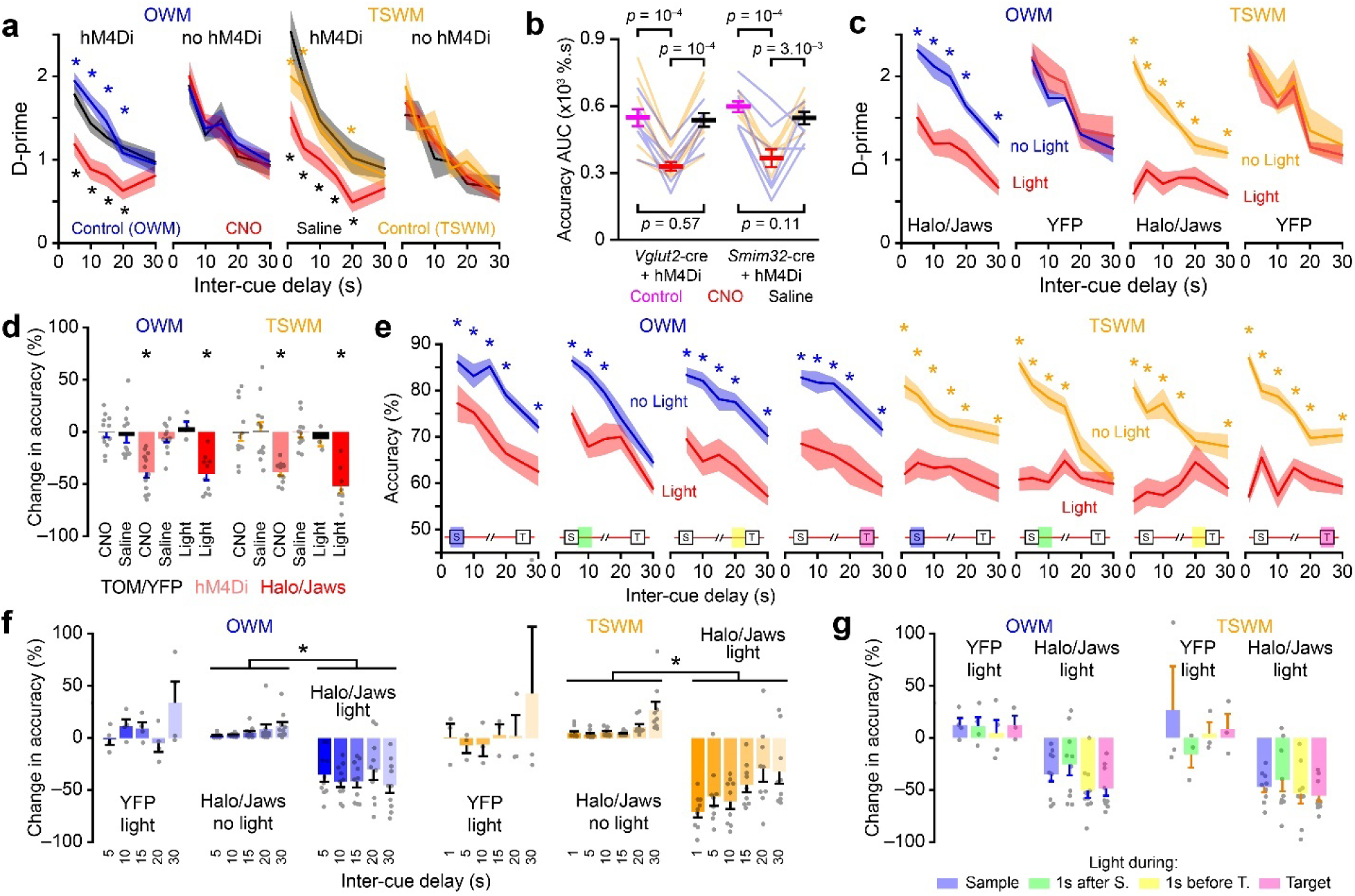
Chemo- and optogenetic inhibition of CLA neurons impair working memory performance. **a**, WM performances calculated from d-prime values and plotted as a function of the inter-cue delay duration for groups of mice either expressing hM4Di in the CLA (OWM: n = 13, TSWM: n = 11) and control mice without hM4Di (n = 12 for OWM and TSWM). Control sessions (no injection), sessions with CNO injection, and sessions with saline injection were performed sequentially in this order (RM ANOVA; post-hoc Fisher’s test comparisons *: P < 0.05, significant comparisons for Control vs. CNO and Saline vs. CNO are color-coded in blue/orange and black, respectively). **b**, WM AUC values quantified in two different transgenic lines targeting the CLA. The lines represent the AUC for individual mice across the different treatments and tasks (OWM [n = 13] and TSWM [n = 11], data are shown in blue and orange, respectively. RM ANOVA with post-hoc Fisher’s test comparisons). **c**, WM performance calculated from d-prime values and averaged across the four stimulation epochs as a function of inter-cue delay duration for mice expressing either Halo/Jaws or YFP in the CLA. Performances for no light and light trials are compared within each group (Number of mice: Halo/Jaws [OWM: n = 10, TSWM: n = 9]; YFP [OWM: n = 4, TSWM: n = 3]. RM ANOVA; post-hoc Fisher’s test comparisons *: P < 0.05, significant comparisons are indicated in either blue or orange). **d**, Change in global response accuracy (relative to control performance, i.e. no CNO or no light conditions) following either chemogenetic or optogenetic CLA inhibition or saline injection in OWM and TSWM tasks. The performance of individual mice is indicated as grey dots. Comparisons were made between CNO and Saline groups using paired t-tests (*: P < 0.05). Comparisons were between YFP control group and Halo/Jaws experimental group using unpaired t-tests (*: P < 0.05). **e**, WM performance for individual stimulation epoch plotted as a function of inter-cue delay duration for mice expressing either Halo/Jaws or YFP in the CLA. Performances for no light and light trials are compared within each group (RM ANOVA; post-hoc Fisher’s test comparisons *: P < 0.05, significant comparisons are indicated in either blue or orange). The schematic of the stimulation epoch is provided within each plot. **f**, Effect of optogenetic inhibition of CLA neurons on WM performance across various inter-cue delays. Accuracy change is calculated for light trials relative to no-light conditions. For Halo/Jaws no-light conditions, performance during the no-light trials of a given stimulation epoch is compared to the averaged performance of the no-light trials across the other three stimulation epochs. (RM ANOVA: *: P < 0.05). **g**, Effect of optogenetic inhibition of CLA neurons on WM performance during different light stimulation epochs for both OWM and TSWM tasks in control and experimental mouse groups (One-Way ANOVA between the different light stimulation epochs P > 0.05 for all comparisons). All data represented as mean ± SEM. See **Supplementary Table 1** for detailed statistics.

**Supplementary Table 1.**
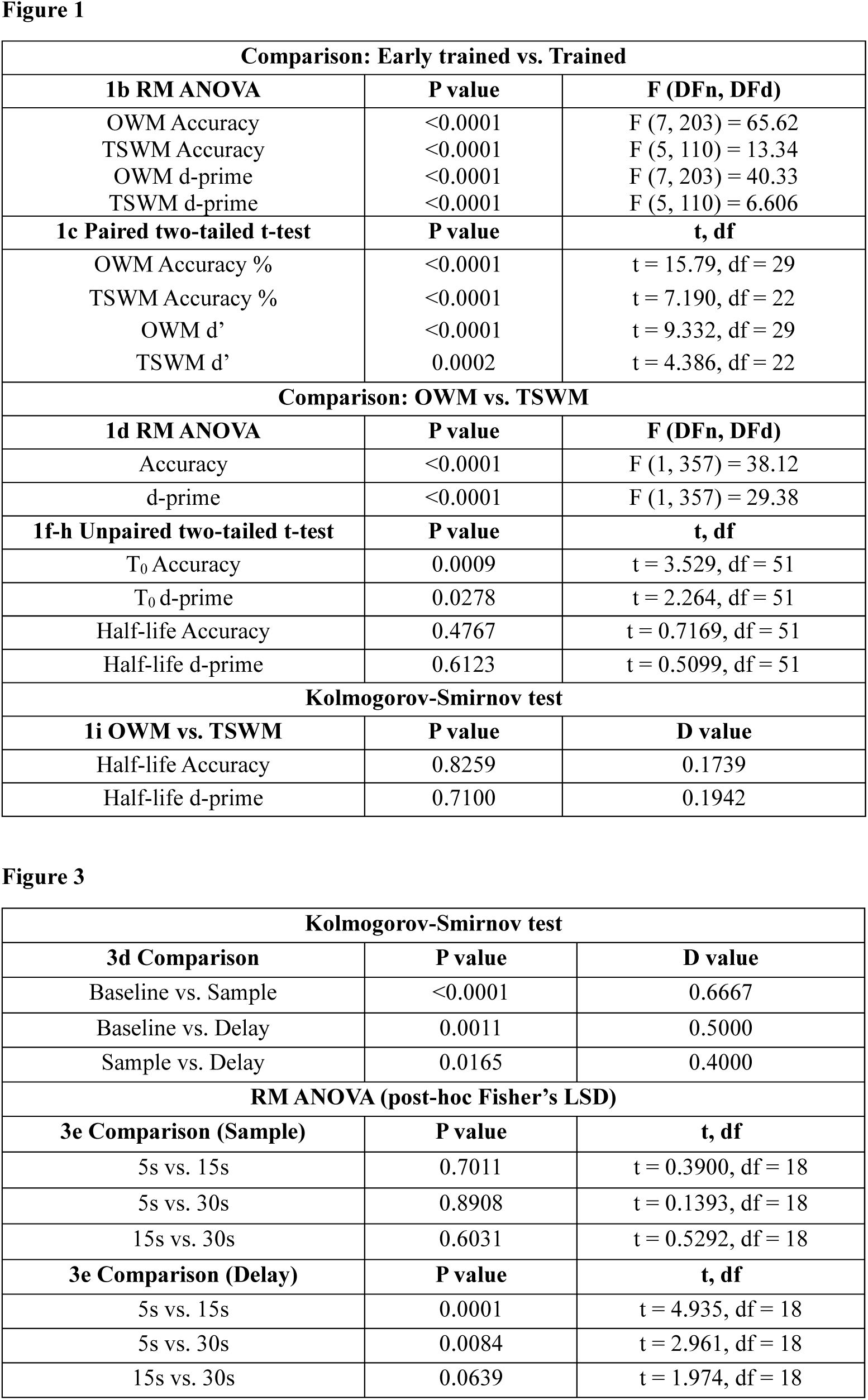
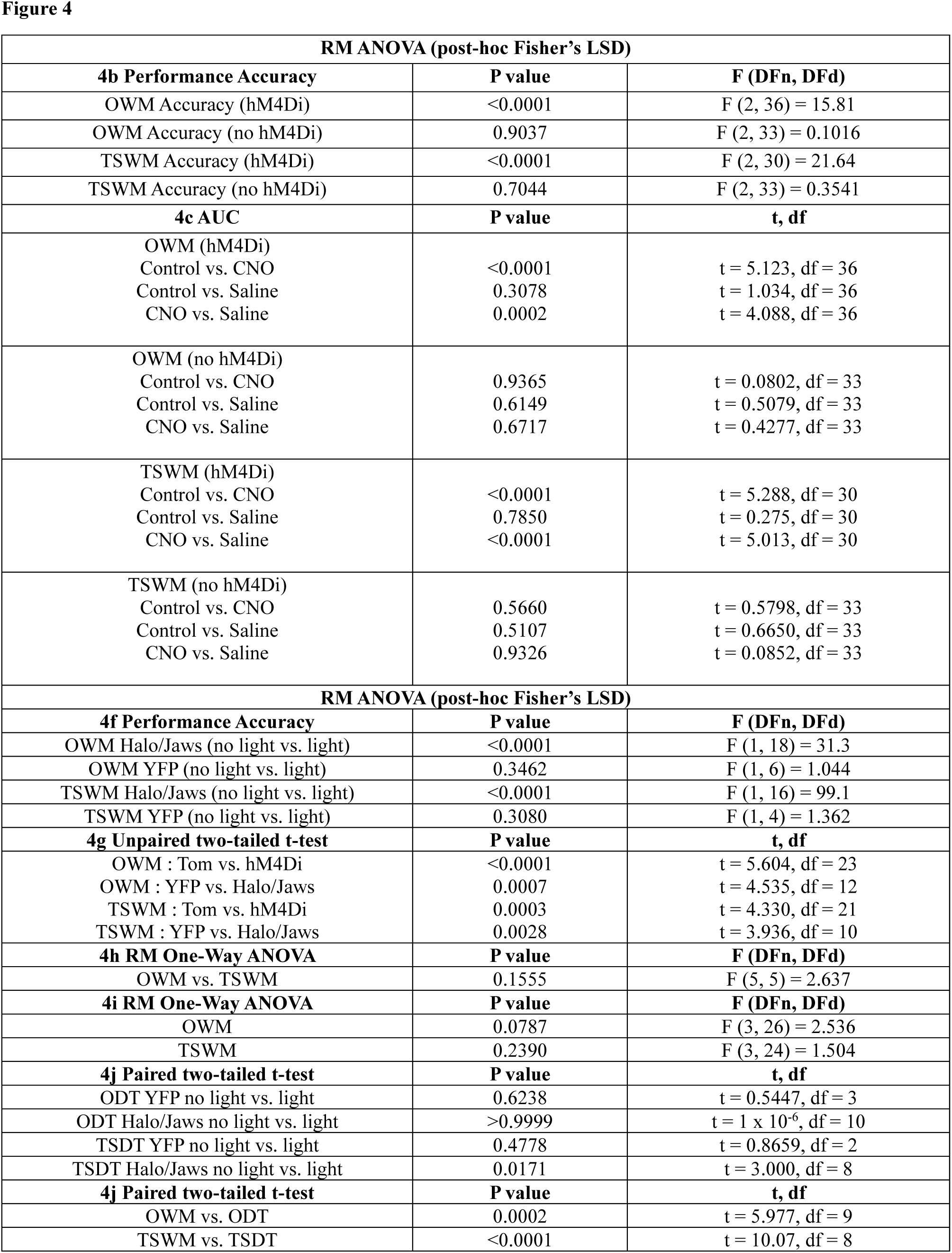
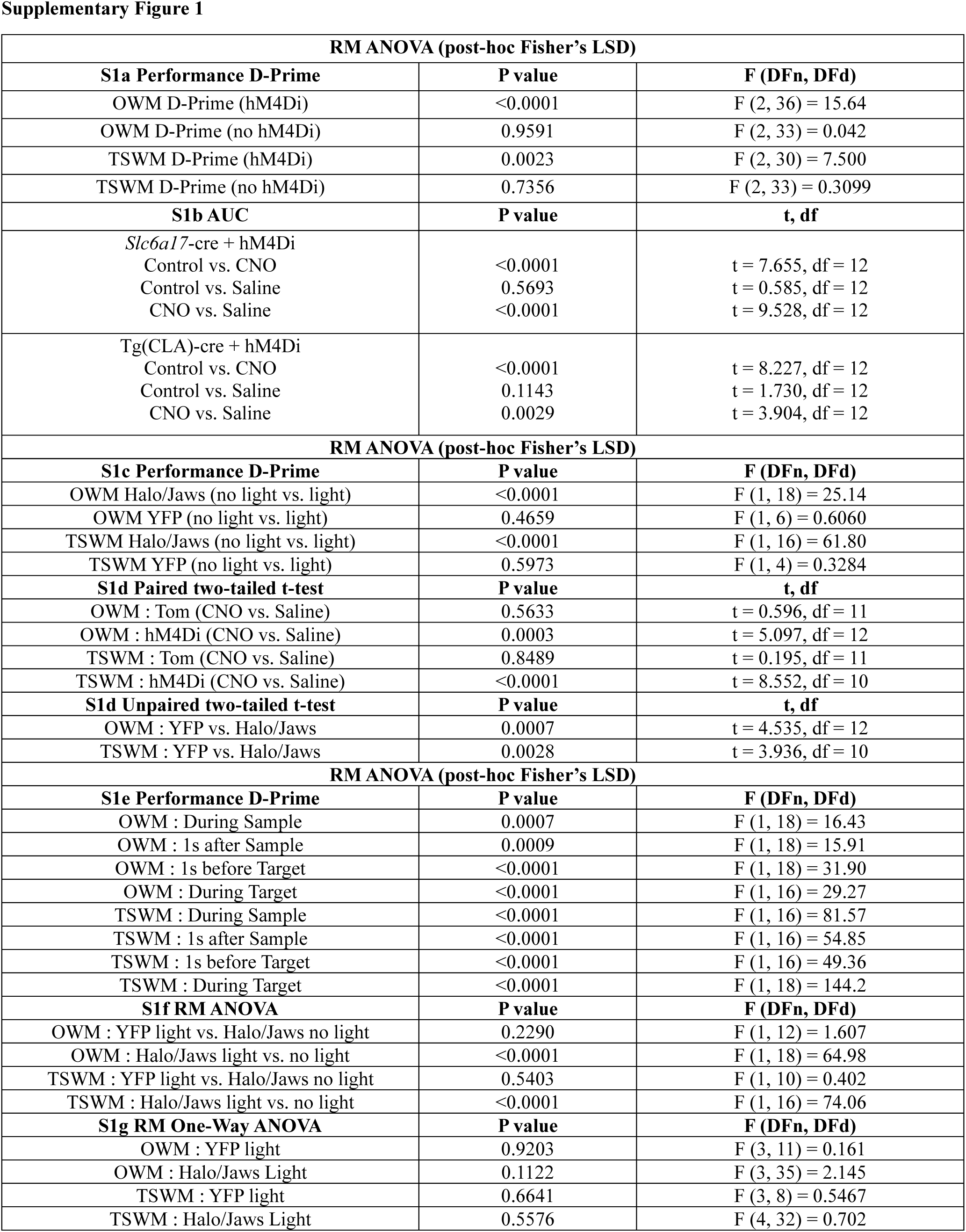
detailed statistics.

## Notes

### Competing Interest Statement

The authors have declared no competing interest.

